# ENT3C: an entropy-based similarity measure for Hi-C and micro-C derived contact matrices

**DOI:** 10.1101/2024.01.30.577923

**Authors:** Xenia Lainscsek, Leila Taher

## Abstract

Hi-C and micro-C sequencing have shed light on the profound importance of 3D genome organization in cellular function by probing 3D contact frequencies across the linear genome. The resulting contact matrices are extremely sparse and susceptible to technical- and sequence-based biases, making their comparison challenging. The development of reliable, robust and efficient methods for quantifying similarity between contact matrix is crucial for investigating variations in the 3D genome organization between different cell types or under different conditions, as well as evaluating experimental reproducibility. We present a novel method, ENT3C, which measures the change in pattern complexity in the vicinity of contact matrix diagonals to quantify their similarity. ENT3C provides a robust, user-friendly Hi-C or micro-C contact matrix similarity metric and a characteristic entropy signal that can be used to gain detailed biological insights into 3D genome organization.

**Availability:** https://github.com/xX3N1A/ENT3C

## Introduction

Chromosome conformation capturing (3C-based) (1) has been pivotal in the discovery of hallmarks of 3D genome organization such as compartments, compartmental domains, topologically associating domains and chromatin loops, that each play distinct roles in regulating gene expression. Specifically, Hi-C (2) and micro-C (3) assays ligate DNA fragments in 3D proximity to one another and use deep sequencing to infer chromatin interaction frequencies. These data are then typically summarized in a *contact matrix* whereby the genome is binned into fixed-sized loci and each entry in the matrix is the estimated interaction between a pair of loci. Contact matrices are typically sparse, and suffer from low signal-to-noise ratios. For example, a low value in the matrix could result from a true lack of contact between the corresponding genomic loci or simply from insufficient sampling. High matrix values may reflect frequent chromatin interactions, or merely spurious ligation products associated with the adopted chromatin fragmentation and ligation protocols (4). In addition, sequence-driven noise in contact matrices arises from nucleotide composition and mappability. Because it helps mitigate stochastic fluctuations or experimental artifacts by emphasizing consistent patterns from which biologically meaningful features like A/B compartments can be determined, contact matrices are often transformed into Pearson correlation matrices where each entry is the correlation coefficient between the corresponding row and column in the contact matrix (2, 5, 6).

Quantifying the similarity between Hi-C and micro-C contact matrices is essential for understanding how the 3D genome organization differs among various cell lines or under different biological conditions. However, the scarcity of the data and presence of intricate artifacts (7, 8) render this task difficult. Traditional statistics such as the Pearson correlation coefficient are not well suited for comparing contact matrices, as they often scores biological replicate contact matrices similarly to unrelated samples (8, 9). To address this limitation, HiCRep (9) implements the *stratum adjusted correlation coefficient (SCC)*, which accounts for the domain structure and distance dependence inherent in contact matrices. Moreover, representing contact matrices as images and networks has facilitated the recognition of their geometric shapes and patterns (10). Similar images will share geometric layouts of local self-similarity (11). Leveraging the analogy, Ardakany et al. (12) developed Selfish, a tool that relies on local structural “self-similarity” to quantify the similarity of two contact matrices. Since, on average, the chromatin interaction frequencies between two genomic loci decrease as their distance increases (2)–resulting in a higher frequency of interactions along the contact matrix diagonal–Selfish restricts its computations to submatrices defined by sliding a window along the contact matrix diagonals. Contact matrices can also be regarded as networks where genomic loci are nodes and the contact frequencies are edges. Thus, GenomeDISCO (13) uses random walks in a network representation of the matrices to smooth them across a range of levels before computing their L1 distance, and HiC-spector (14) quantifies the similarity between contact maps applying spectral graph theory. A gold standard has yet to be established (see (8) for an extensive review).

Our method, which we call “Entropy 3C” (ENT3C), is motivated from both the image processing and network analysis perspectives (10). Like Selfish, our approach to quantifying contact matrix similarity draws upon insights from image processing. The abundant and intertwined features that characterize an image such as its texture, edges, spatial correlations, luminescence and frequency distributions are difficult to disentangle and all contribute to its overall complexity. In particular, the amount of information in an image increases with increasing complexity, which is reflected in higher information entropy. Consistently, entropy has proven valuable in image analysis, aiding in tasks like face detection, character recognition, and parameter determination for augmented reality (15). Not surprisingly, the concept of entropy has also found application in network theory (16). Building on these results, ENT3C detects local changes in the signal near the diagonal of a Hi-C or micro-C contact matrix based on the von Neumann information entropy and recent work concerning entropy quantification of Pearson correlation matrices (17). The von Neumann entropy (18) is a definition of entropy used to describe quantum systems and is equivalent to the classical Shannon entropy under certain conditions. Given a contact matrix, ENT3C computes the entropy of smaller Pearson-transformed submatrices extracted along its main diagonal. We define the similarity between two contact matrices as the Pearson correlation between their entropy signals. We show that our similarity score is robust to both sequencing depth and contact matrix binning resolution and can perfectly distinguish between Hi-C and micro-C contact matrices derived from different cell lines and those derived from biological replicates of the same cell line. We also demonstrate the potential of these entropy signals to investigate intra- and intercellular differences in 3D chromatin interactions.

## Supplementary Note 1: Methods

### A. Concepts of entropy form the basis for ENT3C

ENT3C compares two Hi-C or micro-C contact matrices by calculating an entropy signal **S** along their diagonals. Specifically, ENT3C computes the von Neumann entropy (18), a generalization of the classical Shannon entropy to quantum mechanical systems defined as

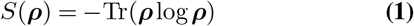

where ***ρ*** is a density matrix representing a quantum mechanical system that is in a mixed state described by an ensemble of pure states with different probabilities and Tr() is the trace. A matrix is a density matrix if and only if it is Hermitian (i.e.,equal to its own complex conjugate), positive semi-definite (i.e., its eigenvalues are greater than or equal to zero), and has unit trace (i.e., its diagonal elements sum to one).

Although, an *N* × *N* contact matrix **A** = (*A*_*kl*_) does not generally satisfy all necessary conditions for a density matrix, its Pearson correlation matrix scaled by 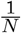 does (17). The Pearson correlation matrix of a contact matrix **A** is a matrix **P** = (*P*_*hk*_) in which each element *p*_*hk*_ corresponds to the Pearson correlation coefficient between the *h*th row and *k*th column of **A** (2).

#### A.1. Entropy signal S

To obtain the entropy signal **S** for a *N*×*N* contact matrix **A** = (*A*_*kl*_), ENT3C extracts a total of *WN* submatrices 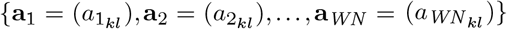 along its diagonal, each of dimensions *n × n* and shifted by *WS* bins. Next, for each submatrix **a**_*i*_, *i* = 1, 2, …, *WN* ENT3C replaces the zeros with the minimum non-zero entry in **a**_*i*_, computes the logarithm of each entry, and transforms the resulting matrix into a Pearson correlation matrix 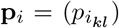 using MATLAB’s function `corrcoef(a,’rows’,’complete’)`. The analogue of the quantum mechanical density matrix is given by 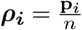, and we can compute the entropy in the equivalent form of Eq.1

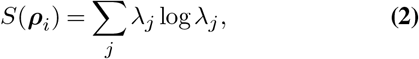

where *λ*_*j*_ is the *j*th eigenvalue of ***ρ***_***i***_ computed with MAT-LAB’s `eig()` function. Values for which *λ* ≤ 0 are ignored. Thus, each contact matrix **A** yields a vector of entropy values **S** = ⟨*S*(***ρ***_1_), *S*(***ρ***_2_),…, *S*(***ρ***_*WN*_)⟩, where *S*(***ρ***_*i*_) represents the entropy of the scaled Pearson correlation matrix of submatrix **a**_*i*_ for *i* = 1, 2, …, *WN* (Figure 1 A).

**Fig. 1.**
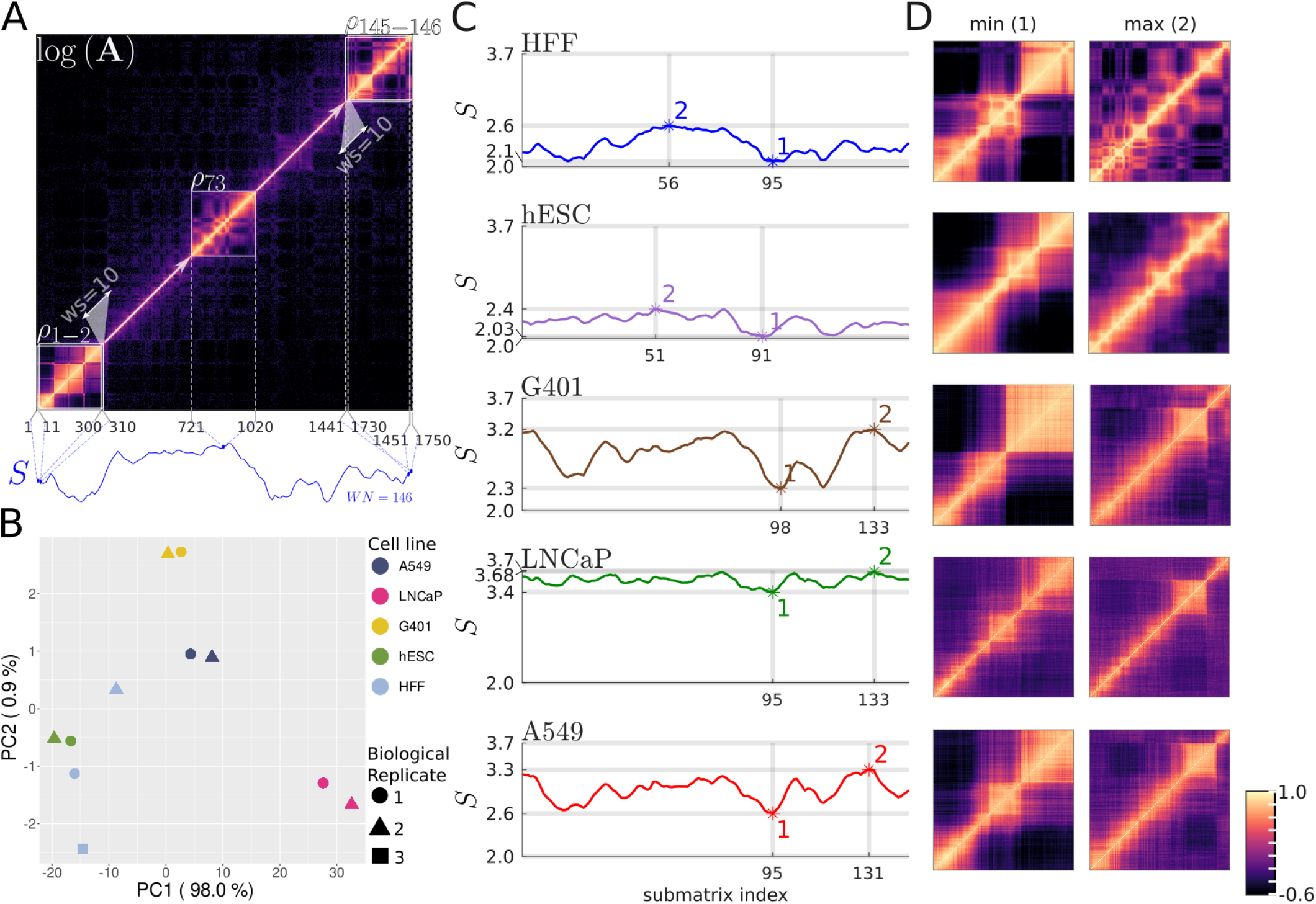
ENT3C computes cell-line specific entropy signals S from the diagonal of a contact matrix and identifies low and high complexity regions. (**A**) Depiction of ENT3C derivation of the entropy signal **S** of a contact matrix **A**. The analysis is exemplified for the pooled BR contact matrix of the HFFc6 cell line for chromosome 14 binned at 40 kb (Methods). ENT3C’s was run with submatrix dimension *n* = 300, window shift *WS* = 10, and maximum number of data points in **S** *WN* _max_ =∞, resulting in *WN* = 146 submatrices along the diagonal of the contact matrix. For subsequent scaled Pearson-transformed submatrices, ***ρ***_*i*_, along the diagonal of log **A**, ENT3C computes the von Neumann entropies *S*(***ρ***_1_), *S*(***ρ***_2_),…, *S*(***ρ***_*WN*_). The resulting signal **S** = ⟨*S*(***ρ***_1_), *S*(***ρ***_2_),…, *S*(***ρ***_*WN*_)⟩ is shown in blue under the matrix. The first two (***ρ***_1−2_), middle (***ρ***_73_), and last two submatrices (***ρ***_145−146_) are shown. (**B**) Principal Component Analysis (PCA) plot of **S** of contact matrices of five cell lines for chromosome 14. Cell lines cluster strongly along PC1 (98% variance explained). The most outlying sample is biological replicate 2 of HFFc6. (**C**) ENT3C entropy signals **S** of pooled BR contact matrices **A** of five cell lines for chromosome 14 was computed using the same parameters as in (A). Minimum (1) and maximum (2) **S** values overlap in some cases. (**D**) Submatrices corresponding to minimum and maximum entropy values in (C) correspond to differences in pattern complexity. Autosome-wide analysis for HFFc6 is given in Supplementary Figure S2.

##### Similarity score Q

ENT3C scores the similarity between two contact matrices **A** and **B** as the Pearson correlation coefficient *r* between the entropy signals **S**_**A**_ and **S**_**B**_. In this context, we refer to *r* as *Q*. When comparing contact matrices, bins are excluded from the calculation if they are empty in any of the matrices of interest.

#### A.3 ENT3C parameters

ENT3C has two key parameters: (1) the submatrix dimension, *n*; and (2) the window shift *WS*. To enable the computation of an informative entropy value, *n* should be at least 50. However, the larger the value of *n*, the smaller the number of points in **S**, and due to insufficient sampling, the corresponding Pearson correlation coefficient might not accurately reflect the true correlation. To make it easier to select a suitable value for *n*, ENT3C’s user can set the *chromosome-split c* instead, which indicates the number of submatrices into which the contact matrix is partitioned. ENT3C then computes *n* as 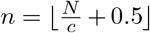, where ⌊⌋ is the floor function.

Additionally, the maximum number of data points in **S** can be controlled to save computation time by setting a *WN* _max_. Note that the number of data points *WN* in **S** is implicitly defined by the combination of *n* and *WS*. If *WN > WN* _max_, *WS* is increased such that *WN* is at most *WN* _max_.

Default ENT3C parameters are *c* = 7, *WS* = 1, and *WN* _max_ = 1000.

### B. Hi-C and micro-C data sets

Processed Hi-C data for the A549, G401, and LNCaP cell lines were obtained from the ENCODE Portal (https://www.encodeproject.org/, last accessed in November, 2023). Specifically, we downloaded the *BAM* files (ENCODE4 v1.10.0 GRCh38) for each of the available replicates. BAM files were transformed into pairs files using the `pairtools parse` function (Supplementary Table S1). These data represent a subset of the data assessed in a recent review on contact matrix similarity metrics (8) and were thus chosen for comparibility.

To also include newer data sets, processed high-resolution micro-C data sets of the H1-hESC and HFFc6 cell lines (3) were obtained from the 4D Nucleome Data Portal (https://data.4dnucleome.org/, last accessed in November,2023). We downloaded *contact list-replicate (pairs)* files for the two H1-hESC and three HFFc6 replicates (Supplementary Table S2).

#### B.1 Pooled technical and biological replicates

The pairs files of the technical replicates of the same micro-C BR (Supplementary Table S2) were merged to create a pairs file for each BR using *pairtools*’s (v1.0.2) `merge` function (19). Similarly, the pairs files of the same cell line were merged to create a pairs file of pooled BRs for each cell line.

#### B.2 Binning and generating contact matrices

Pairs were aggregated into binned contact matrices and stored in HDF5 `cooler` format using cooler’s (v0.8.11) (20) `cload pairs` function.

#### B.3. Downsampling pairs

When specified, pairs files were downsampled with *pairtool*’s (v1.0.2) (19) `sample` function.

### C. Identifying high and low complexity regions within a contact matrix

To identify the submatrices corresponding to the highest and lowest entropy values in **S**, ENT3C was run on 40 kb-binned pooled BR contact matrices of chromosome 14 with parameters *n* = 300, *WS* = 10 and *WN* _max_ = 1000 for all cell lines. The same analysis was performed for all autosomes for the HFFc6 cell line.

### D. Quantifying contact matrix similarity with ENT3C

#### D.1 Assessing average performance of the similarity metric

We evaluated ENT3C’s performance by its ability to distinguish contact matrices obtained from biological replicates of the same cell line (BRs) from those derived from different cell lines (NRs).

Specifically, for cell line *i*, we first calculated the similarity score *Q* between every pair of replicates for each chromosome, then averaged *Q* across all chromosomes, and then across all pairs of replicates. This is further referred to as 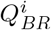. The average of 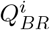 across all cell lines is denoted 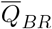.

Analogously, for each pair of cell lines *i* and *j*, we first calculated the similarity score *Q* for each chromosome between every possible combination of replicate pairs available for the cell line *i* and *j*. (Supplementary Table S1-S2). Then we averaged *Q* across all chromosomes, and then across all pairs of replicates. This is further referred to as 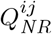. The average of 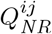 across all cell lines is denoted as 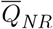.

Furthermore, for cell line *i*, we computed a separating margin *d*_*i*_ as the distance between 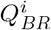 and the median of 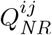 across all cell lines *j* with *j* ≠ *i*. The average of *d*_*i*_ across all cell lines is denoted 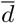.

#### D.2 Testing the sensitivity of ENT3C’s parameters n and WS

We tested the sensitivity of ENT3C’s parameters *n*, the submatrix dimension, and *WS*, the window shift, on 40 kb-binned contact matrices of chromosome 14. Matrices derived from downsampled pairs files (30 million interactions) were used to ensure comparability. First, we fixed *WN*_*max*_ = ∞, *WS* = 1 and let *n* ∈ {50, 263, 476, 689, 902, 1114, 1327, 1540, 1743}. Secondly, we fixed *n* = 300 and varied *WS* ∈ {1, 63, 126, 188, 251, 313, 375, 438, 500}.

#### D.3 Benchmarking ENT3C in regard to sequencing depth

We tested ENT3C’s sensitivity to the sparsity of the contact matrix by computing 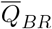 and 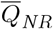 on 40 kb-binned contact matrices derived from pairs files randomly downsampled to contain 5, 10, 20, 25, 30, 60, 120, 240, 400, and 800 million interactions, provided the original pairs file contained a sufficient number of interactions (Supplementary Table S3-S4). For this analysis, *WS* = 1 and *WN* _max_ = 1000; the chromosome-split was fixed to *c* = 7 to account for the chromosome-dependent variation in the dimensions of the contact matrices.

#### D.4 Benchmarking ENT3C in regard to binning resolution

We tested ENT3C’s sensitivity to binning resolution by computing 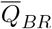 and 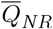 on contact matrices derived from intact pairs files at 8 different binning resolutions: 5, 10, 25, 40, 50, 100, and 500 kb. Specifically, we first generated 5 kb-binned contact matrices from intact `pairs` files and then recursively coarsened them to lower resolutions with `cooler`’s (v0.8.11) (20) `zoomify` function. To obtain a sufficiently large number of data points *WN* in **S**, we set *c* depending on the binning resolution: *c* = 150 for 5 kb, *c* = 100 for 10 kb, *c* = 25 for 25 kb, *c* = 7 for 40 kb and 50 kb, *c* = 6 for 100 kb, *c* = 3 for 500 kb, and *c* = 2 for 1 Mb. Further, we set *WS* = 1 and *WN* _max_ = 2000.

### E. Comparisons to state-of-the art methods

HiC-Spector (`run_reproducibility_v1.py`) (14), QuASARRep (21), GenomeDISCO (v1.0.0) (13), and HiCRep (v1.6.0) (14) were run using the wrapper in the `3DChromatin_ReplicateQC` github repository (version 1.12.2). All tools were run following the author’s recommendations to the best of our abilities, using the parameters file provided by `3DChromatin_ReplicateQC` where applicable. We converted the cool files of each contact matrix into the input format required by `3DChromatin_ReplicateQC` using `HiCExplorer`’s (v3.7.2) (22) `hicConvertFormat` function with options “`-inputFormat cool -outputFormat ginteractions`” and subsequently rearranging the columns. As HiC-Spector, QuASAR and GenomeDisco implement internal normalization algorithms, we called `hicConvertFormat` with the option “`-load_raw_values`” to extract unbalanced matrices from the cooler containers. The balanced matrices from the output of the cooler `balance function` with option “`-max-iters 300`” were used as input for HiCRep. Selfish (12) (MATLAB implementation committed on Feb 15, 2019) accepts contact matrices in Rao (hic) format and its two parameters *k* and *c* are determined empirically. Parameter *k* is directly proportional to the blocksize of the submatrices which are slid over the diagonal and must be chosen such that the resulting submatrices enclose structures likely to be preserved between cell lines, such as TADs; *c* depends on the Hi-C protocol. We fixed *k* = 100 as the authors did in their original publication, since our contact matrices were of the same binning resolution. As Hi-C and micro-C protocols are different, we initially set *c* = 10 and iterated for *c* = *c* − 1 to find the largest integer where similarity score for BRs reached 0.9 as suggested by the authors. ENT3C was run with default parameters (*c* = 7, *WS* = 1 and *WN* _max_ = 1000).

We compared similarity scores of all methods for (1) intact contact matrices, and (2) contact matrices derived from pairs files downsampled to 30 million interactions. Performance of each method was determined by their respective similarity scores averaged across BR and NR contact matrices (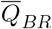 and 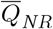, respectively), and separating margin 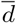.

### F. Identifying high and low complexity regions within and between two contact matrices

To identify the most similar genome-wide regions between HFFc6 and H1-HESC (pooled BRs) we fit a linear model using R’s `lm` function to the entropy signals of their contact matrices binned at 5, 10, 25, 40 and 50 kb resolution. Then we selected the regions closest to the linear fit (below the 0.3% quantile) and extracted the Ensembl IDs of protein-coding genes with transcription start sites falling within these regions. ENT3C was run with chromosome-split *c* = 150 for 5 kb, *c* = 100 for 10 kb, *c* = 25 for 25 kb, *c* = 7 for 40 kb and 50 kb, *c* = 6 for 100 kb, *c* = 3 for 500 kb, *c* = 2 for 1 Mb. The window shift *WS* was set to *WS* = 1 and *WN* _max_ = 2000.

#### F.1 Functional enrichment analysis

Annotation data were obtained from BioMart (GRCh38.p14). Enrichment for Gene Ontology (23) terms describing *biological processes* was statistically assessed with `clusterProfiler`’s (24) `compareCluster` (v.4.10) function with parameter ‘`qvalueCutoff=0.05`’.

## Supplementary Note 2: Results

### Variations in local entropy patterns indicate changes in 3D genome organization

Within a cell, regions in the genome characterized by complex 3D organization will be reflected by more intricate patterns in their Hi-C or micro-C contact matrices. The degree of complexity of these patterns can be quantified by entropy. Given an *N*× *N* contact matrix, ENT3C generates an entropy signal **S** by calculating the von Neumann entropy (18) of *n*×*n* scaled Pearson-transformed submatrices along the diagonal, each shifted by *WS* (Figure 1A, Supplementary Figure S1; Methods). Supporting our hypothesis that variations in local entropy patterns indicate changes in cell-line specific 3D organization, **S** resulting from biological replicates (BRs) of the same cell line tended to be more clustered along principal component 1 than those of different cell lines (non-replicates: NRs) (Methods; Figure 1B).

To illustrate how **S** can capture differences in the complexity of different regions of a contact matrix, we identified the regions of minimum and maximum entropy values of chromosome 14 for pooled BRs of five different cell lines (Methods; Figure 1C). Indeed, we found that higher entropy values reflected more intricate patterns in the corresponding contact matrices, particularly in HFFc6 (Figure 1D, Supplementary Figure S2). Interestingly, we identified nearly the same low complexity region for each cell line, suggesting this region’s transcriptional landscape is conserved between cell lines. Moreover, HFFc6 and H1-hESC displayed similar high complexity regions, as did G401, LNCaP, and A549.

### B. ENT3C’s similarity score accurately captures contact matrix similarity

ENT3C quantifies the similarity between two contact matrices **A** and **B** by computing the Pearson correlation coefficient between their entropy signals **S**_**A**_ and **S**_**B**_, to which we further refer as the *similarity score Q*. Remarkably, for fixed parameters *n* = 300, *WS* = 1 and *WN* _max_ =∞, ENT3C effectively distinguished between (intact) 40 kb-binned contact matrices derived from biological replicates of the same cell line (BR) and those derived from different cell lines (NR), with average similarity scores of 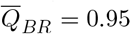 and 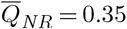, and average separating margin 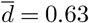 (Methods; Supplementary Figure S3 A). Moreover, its discrimination power remained stable (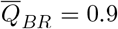 and 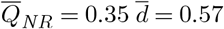) upon downsampling to 30 million interactions to account for variations in library size (Methods; Supplementary Figure S3 B).

By only analyzing submatrices near the diagonal, ENT3C captures signals originating from dominant patterns in the contact matrix. This renders it notably insensitive to sampling depth and binning resolution. To verify robustness to the sequencing depth of a Hi-C or micro-C experiment, we applied ENT3C to contact matrices comprising various numbers of interactions (Methods; Supplementary Table S4), and found that ENT3C consistently distinguished between BR and NR contact matrices (Figure 2 B and Supplementary Figure S4 B). Similarly, we tested ENT3C’s robustness to the binning resolution of the contact matrix for matrices generated at 8 binning resolutions ranging from 10 kbp to 1 Mbp (Methods). Once more, ENT3C consistently distinguished between BR and NR contact matrices (Figure 2 A and Supplementary Figure S4 A).

**Fig. 2.**
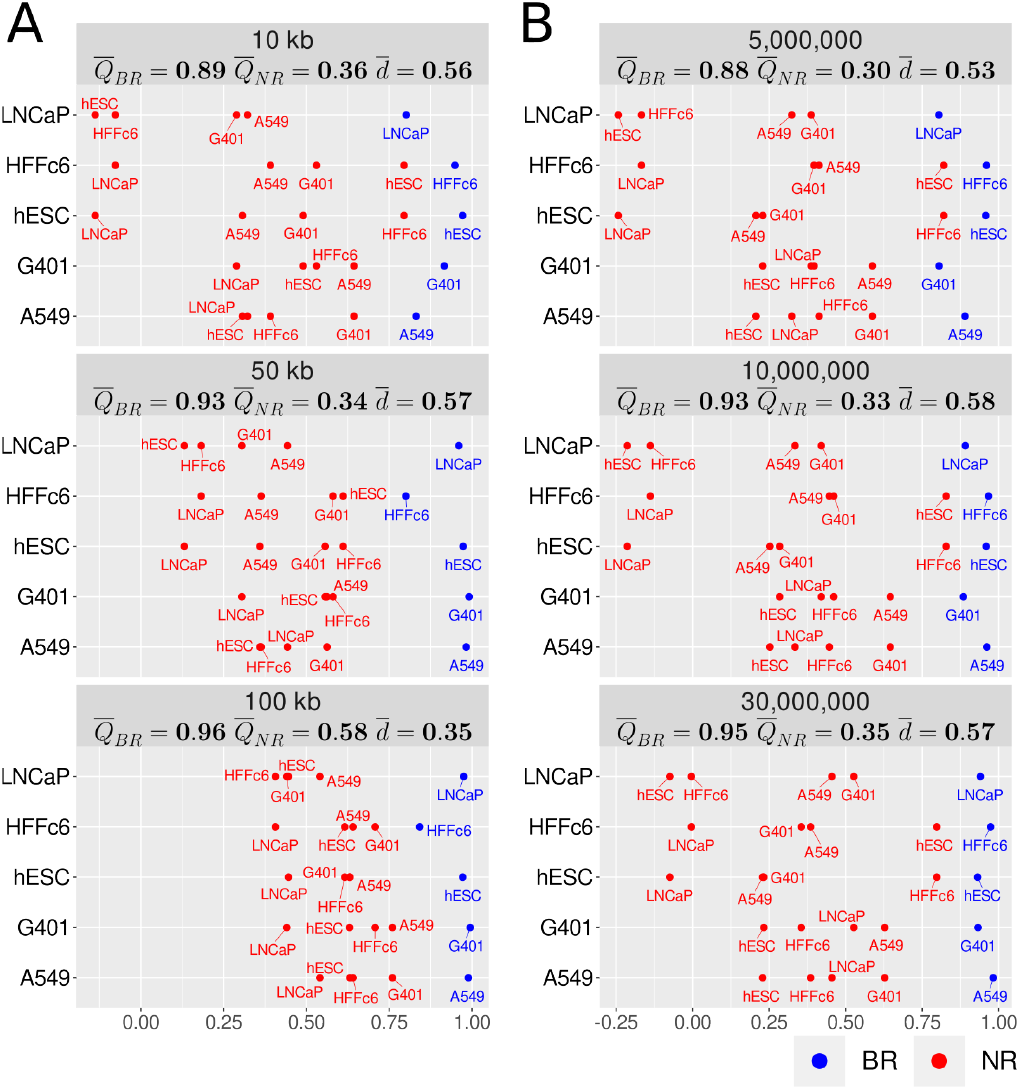
ENT3C is insensitive to binning resolution and sequencing depth. For each cell line *i* indicated on the *y*-axis, a blue dot represents the average ENT3C similarity score across all chromosomes and biological replicate pairs (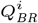 Methods) and a red dot represents the average ENT3C similarity score with cell line *j* (indicated on the label) computed across all chromosomes and replicate pairs (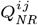 ; Methods) for **(A)** intact contact matrices binned at 10, 50 and 100 kb resolutions, and **(B)** 40 kb contact matrices generated from pairs files downsampled to 5, 10, and 30 million interactions. Panel labels show the averages values across all cell lines or pairs of cell lines (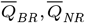), and the average separating margins (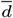). ENT3C was run with parameters *c* = 7, *WS* = 1, and *WN* _max_ = 1000. Other resolutions and depths are given in Supplementary Figure S4.

To further show that ENT3C only marginally depends on the choice of its two key parameters (Methods), we computed 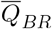 and 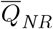 for various values for the submatrix dimension, *n*, and window shift, *WS* (Supplementary Figures S5 A and B). Although the separating margins 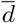 (Methods) did decrease with decreasing *n* due to the loss of information, ENT3C’s similarity measure still correctly distinguished BR from NRs even for extremely small matrix dimensions (for *n* = 50, 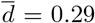 Supplementary Figure S5 A). For the largest submatrix dimension (*n* = 1743), **S** contained merely *WN* = 9 data points, rendering the Pearson correlation coefficient between two signals a poor and unreliable similarity measure.

Regarding the window shift (*WS*) parameter, ENT3C failed for *WS* ≥ 313 (*d* = 0.16), which was very close to the submatrix dimension *n* chosen for the analysis (*n* = 300; Supplementary Figure S5 B). This can be explained by the omission of presumably relevant segments along the diagonal of the contact matrix.

### C. ENT3C competes well with other methods quantifying 3C contact matrix similarity

ENT3C’s performance is similar to that of other state-of-the-art methods. Moreover, ENT3C shows a wider separating margin *d* between BR and NR similarity scores (Methods). To ensure comparability, we performed comparisons both on the intact contact matrices as well as on contact matrices in which the interaction pairs file was downsampled to 30 million genome-wide interactions (Methods). For the intact contact matrices ENT3C, HiCRep and Selfish achieved perfect separation between BR and NR samples, whereby ENT3C’s similarity scores displayed the highest average separating margin between BR and NR samples (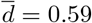) (Supplementary Figure S6; Table 1 A). We observed a similar trend when the contact matrices were derived from downsampled pairs files. ENT3C resulted in the highest average separating margin between BR and NR samples (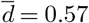) (Figure 3; Table 1 B).

**Table 1.** Comparison of ENT3C to other state-of-the-art methods for quantifying Hi-C and micro-C contact matrix similarity. 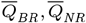, and 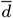 (Methods) values correspond to Figure 1 and Supplementary Figure S6. **A**) intact contact matrices. **B**) contact matrices derived from downsampled (30 million interactions) pairs files.

|  | ENT3C | Genome DISCO | HiC-Spector | HiCRep | QuASAR-Rep | Selfish |
| --- | --- | --- | --- | --- | --- | --- |
| **A) Intact** |  |  |  |  |  |  |
| $\overline{Q}_{BR}$ | 0.93 | 0.80 | 0.32 | 0.95 | 0.70 | 0.93 |
| $\overline{Q}_{NR}$ | 0.34 | 0.43 | 0.17 | 0.59 | 0.48 | 0.68 |
| $\bar{d}$ | 0.59 | 0.49 | 0.06 | 0.40 | 0.24 | 0.27 |
| **B) Downsampled** |  |  |  |  |  |  |
| $\overline{Q}_{BR}$ | 0.95 | 0.71 | 0.29 | 0.88 | 0.26 | 0.93 |
| $\overline{Q}_{NR}$ | 0.35 | 0.39 | 0.16 | 0.53 | 0.16 | 0.68 |
| $\bar{d}$ | 0.57 | 0.50 | 0.06 | 0.39 | 0.01 | 0.26 |

**Fig. 3.**
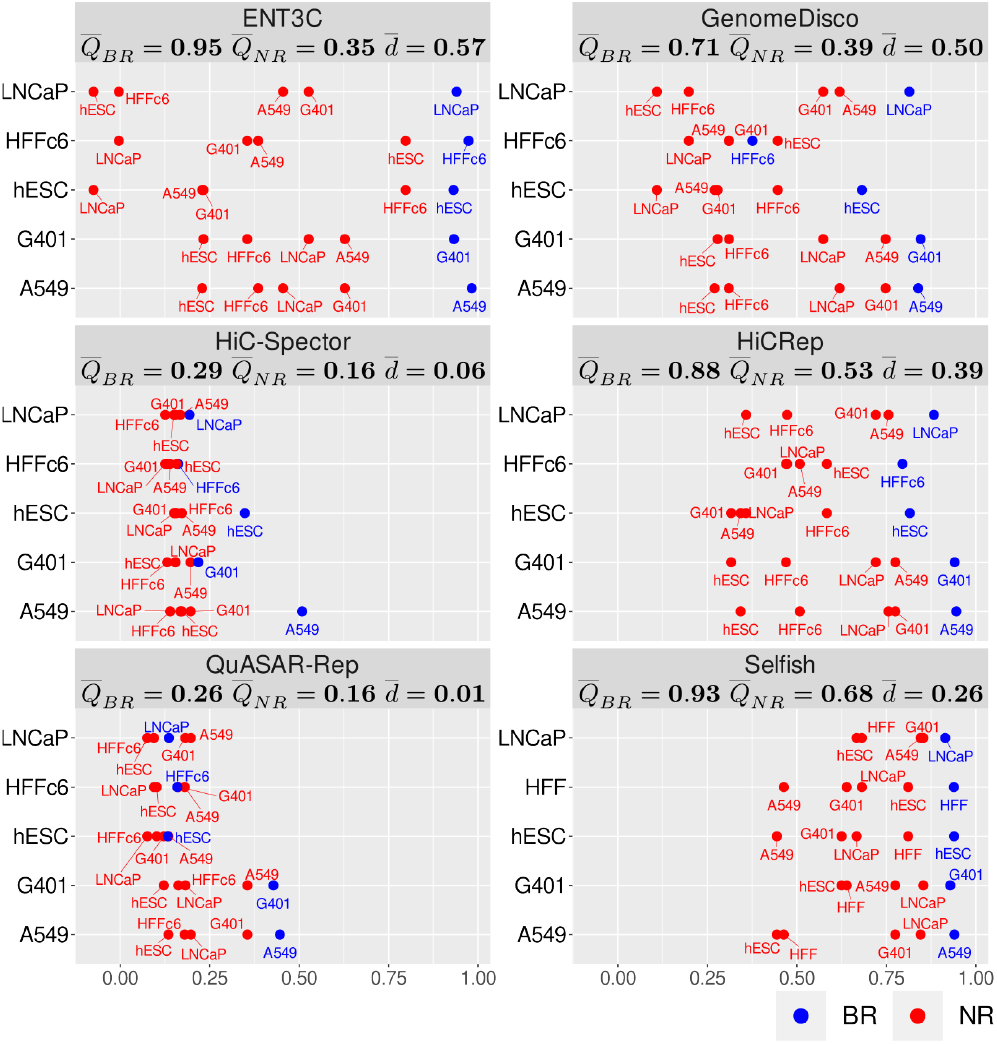
ENT3C competes well with other methods quantifying Hi-C or micro-C contact matrix similarity. Each panel represents a Method (ENT3C, GenomeDISCO, HiC-Spector, HiCRep, QuASAR and Selfish) and each dot represents the average similarity scores 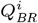 and 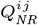 as in Figure 2 (Methods). 40-kb binned contact matrices derived from downsampled pairs files (30 million interactions) were used to ensure comparability. Panel labels show the averages values across all cell lines or pairs of cell lines (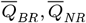), and the average separating margins (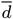). ENT3C was run with parameters *c* = 7, *WS* = 1, and *WN* _max_ = 1000.

Remarkably, the similarity scores of ENT3C’s, HiCRep, Selfish and GenomeDISCO agreed in many cell-line specific differences. For example, when examining the LNCaP cell line, differences to the HFFc6 and H1-hESC cell lines were more pronounced than those between G401 and A549. (First rows ENT3C, HiCRep, GenomeDisco, HiCRep and Selfish in Figure 3 and Supplementary Figure S6). Similar trends were evident for other cell types as well. Like ENT3C, Selfish’s similarity scores remained especially robust regardless of downsampling.

Finally, it is noteworthy that ENT3C, HiCRep and Selfish similarity metrics displayed chromosomal dependency (Supplementary Figure S7). This effect is highly evident, for example, on chromosome 19, which has many unique features such as having the highest GC content of all chromosomes (41%) and more than double the genome-wide average gene density (25).

### D. ENT3C uses entropy to reveal regions of varying complexity between cells

To demonstrate how ENT3C can be used to compare the 3D chromatin organization of different cell lines, we identified regions with the most similar entropy values between the HFFc6 and H1-HESC cell lines at resolutions between 5-50 kb (Supplementary Figure S8-S9, Methods) At low resolutions, the most similar regions identified by ENT3C comprised genes associated with immune response (Methods), which are known to be expressed in nearly all nucleated cells and play a vital role in adaptive immune response (e.g., (26)). At high resolutions, the most similar regions identified by ENT3C were associated with biological processes involved in structure and organization (nucleosome assembly, intermediate filament organization) or DNA-templated transcription initiation, which are also vital for a majority of cells (27). Interestingly, many processes were resolution-specific, which is consistent with distinct gene regulation mechanisms being captured at different resolutions (28).

## Supplementary Note 3: Discussion

Most state-of-the-art methods for quantifying the similarity between two contact matrices apply dimensionality-reducing transformations aimed at modeling and reducing noise and artifacts before computing a similarity score (8). Similarity scores are calculated as the correlation of valid values from the transformed matrices (21), the weighted sum of distance-stratified Pearson correlation coefficients (14), the difference in random walks in a network representation of the matrices (8, 9, 13, 21), or by spectral decomposition of a network representation of the matrices (14), among other methods. ENT3C uses a radically different approach based on the von Neumann entropy (18). Contact matrices can be interpreted among others as images or networks, both of which have been analyzed in other contexts by various definitions of entropy. Moreover, 3D chromatin organization as well as 3C contact matrices have long been associated with fractals (e.g. (29– 33)) further motivating the application of entropy measures to their analysis (34, 35). The von Neumann entropy is under certain conditions identical to the classical Shannon entropy, but is more suitable for analyzing contact matrices because it considers both the positions and the distribution of values in a matrix.

Alternative tools were run with default settings and might perform better upon parameter optimization. Nonetheless, ENT3C achieved the largest average separating margin between contact matrices derived from biological replicates of the same cell line (BR) and those derived from different cell lines (NR) when compared to other state-of-the-art methods.

This result is relevant because a similarity score with a wide separating margin can not only make a binary distinction between BR and NR derived contact matrices, but yield a continuous representation of the 3D organization across diverse cell lines. Interestingly, we observed that method performance tended to be chromosomal-dependent, indicating that the various approaches –and potentially those that we may have inadvertently omitted given the rapidly evolving nature of this field– pick up on different signals in the contact matrices and could be combined into an even stronger similarity metric.

ENT3C disregards any interactions beyond a submatrix dimension from the diagonal. While we might overlook important cell-line differences in the process, the sparsity of long range interactions in current Hi-C and micro-C datasets and gained computational efficiency, justifies this strategy. As data quality increases, ENT3C could easily be modified to take into account longer range interactions. In general, ENT3C’s performance is only marginally dependent on the dimension of its input parameters, i.e., the dimension of the submatrices *n* and the size of the window shift *WS*. Naturally, as fewer data points are generated, a gradual decrease in the information contained in the signal becomes apparent and the similarity metric breaks down. By default, ENT3C automatically sets a suitable value of *n* based on the dimension of the entire contact matrix. The computationally most demanding step of ENT3C is computing the Pearson matrix and eigenvalues; nonetheless after loading the contact matrix and excluding empty bins, computing **S** for chromosome 1 of HFF (biological replicate 1) binned at 40 kb (1753 bins) takes merely ∼10s using parameters *n* = 300, *WS* = 1 and *WN* _max_ = ∞ (MATLAB version 9.14.0.2337262 (R2023a) Update 5 on an AMD^©^ Ryzen 9 3900×12-core processor ×24; Ubuntu 20.04.6 LTS). ENT3C’s high performance and computational efficiency also renders it ideal for training and testing computational models which predict contact matrices.

By tracking subtle differences in the complexity of local patterns along the diagonals of two contact matrices, ENT3C is not only able to distinguish BRs from NRs, but can also be used to investigate the 3D genome organization of one or more cell types in more detail to reveal potentially biologically significant features. The entropy values are difficult to interpret, but the upper value of the von Neumann entropy is confined by the (sub)matrix dimension, and we observed clear differences in the complexity pattern of the submatrices yielding the highest and lowest value in **S**. Likewise, regions exhibiting most similar entropy values between two cell lines were associated with specific biological processes which varied depending on the resolution of the contact matrices, confirming that distinct regulatory mechanisms are captured by different resolutions.

Entropy has long been established beyond its original definitions in thermodynamics and statistical mechanics, across a diverse array of fields. We have demonstrated its potential in 3D genomics by applying it to the challenging problem of quantifying contact matrix similarity. ENT3C provides an efficient and robust similarity measure to assess the reproducibility of Hi-C and micro-C experiments, crucial for coming to cell-type- and condition-dependent conclusions.

## Supplementary Note 4: Data availability

ENT3C is publicly available in MATLAB and Julia on GitHub (https://github.com/xX3N1A/ENT3C).

## Notes

https://github.com/xX3N1A/ENT3C

### Competing Interest Statement

The authors have declared no competing interest.

### Summary of Updates

All Figures updated; Additional results Figure 2; Clarified Methods (mathematical notations, use of experimental data sets); Supplemental files updated

