## Supplementary Material for "ENT3C: an entropy-based similarity measure for Hi-C and micro-C derived contact matrices"

### Figures

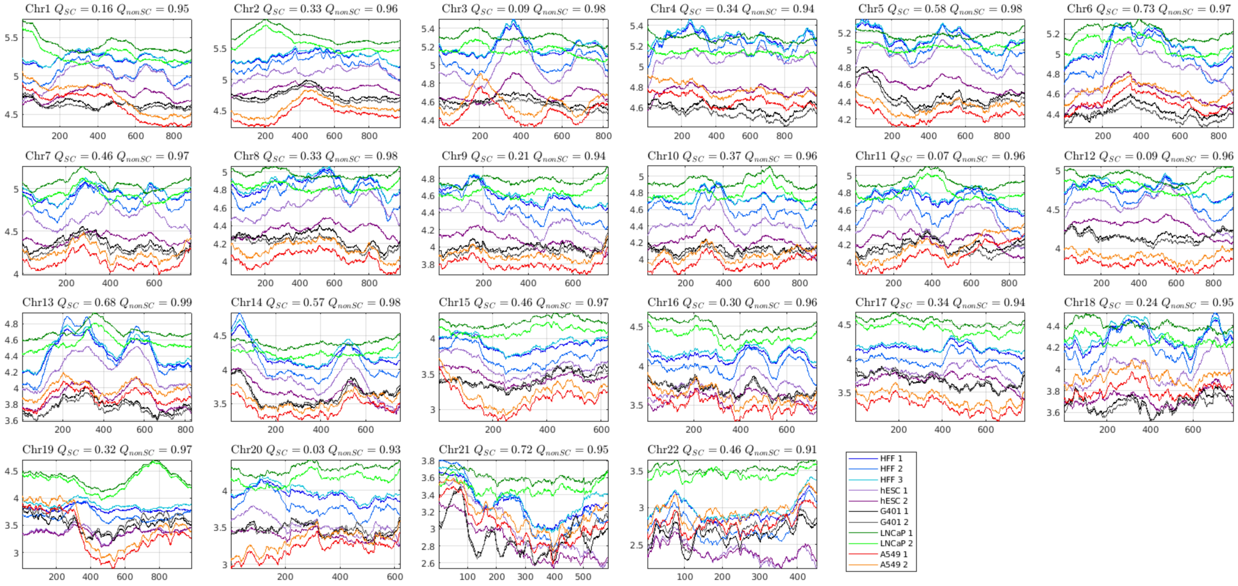


Figure S1: **ENT3C uses the Pearson correlation of entropy signals S to define contact matrix similarity. S** is shown for 40 kb-binned contact matrices from pairs files downsampled to 30 million interactions in various cell lines. Titles indicate ENT3C similarities $Q$ (Methods) between contact matrices derived from biological replicates of the same cell line (BR) and non-replicates derived from different cell lines (NR). BRs are represented in similar color schemes (HFFc6 (blues), H1-hESC (purples), G401 (grays), LNCaP (greens), A549 (reds)).

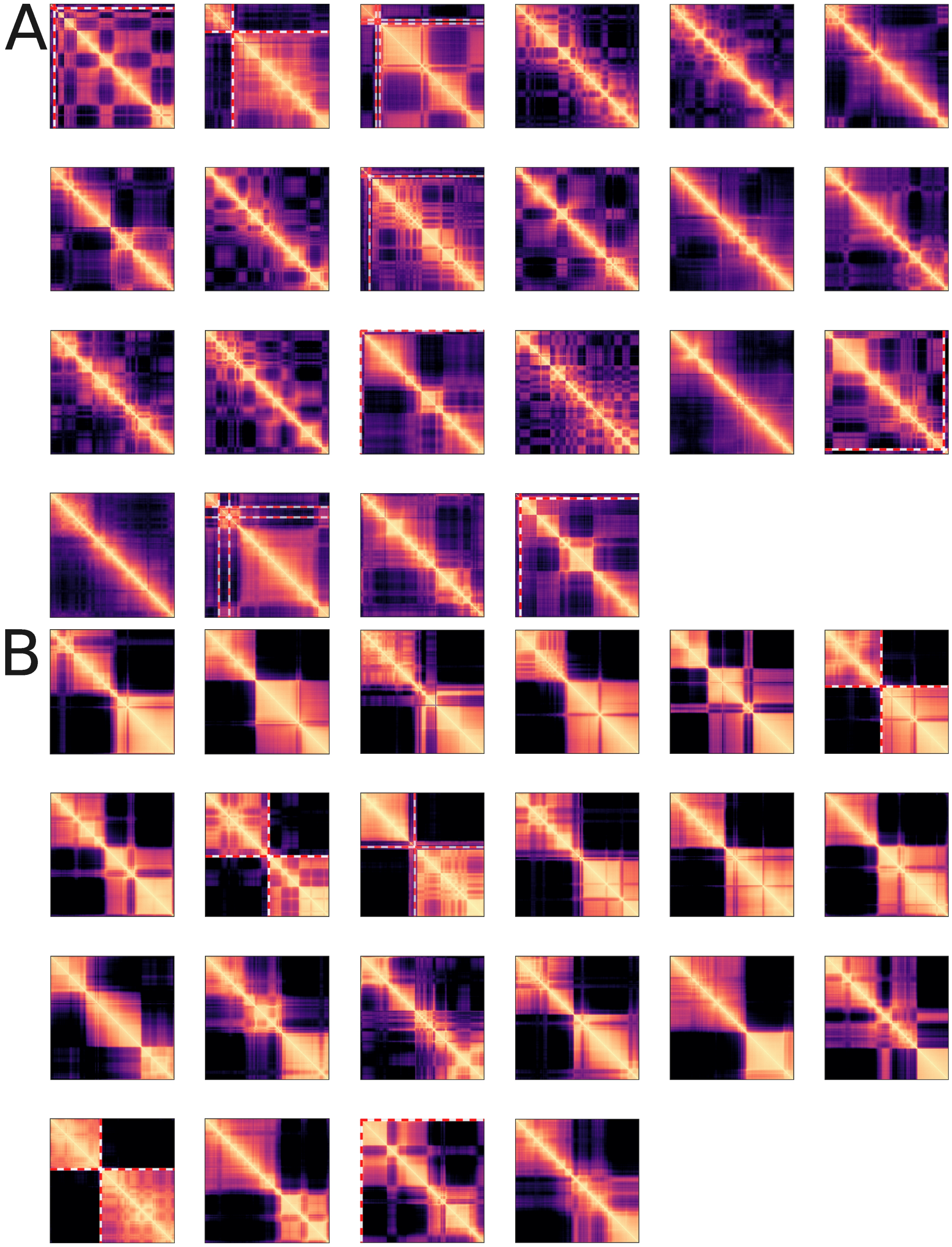


Figure S2: **Highest and lowest entropy submatrices of each chromosome correspond to higher and lower pattern complexity respectively.** Submatrices corresponding to maximum **(A)** and minimum **(B)** entropy values of each chromosome of 40 kb binned HFFc6 contact matrices (pooled biological replicates). ENT3C parameters: submatrix dimension $n = 300$, window shift $WS = 10$, maximum number of matrices evaluated $WN_{\max} = \infty$. White and red stripes indicate centromere regions.

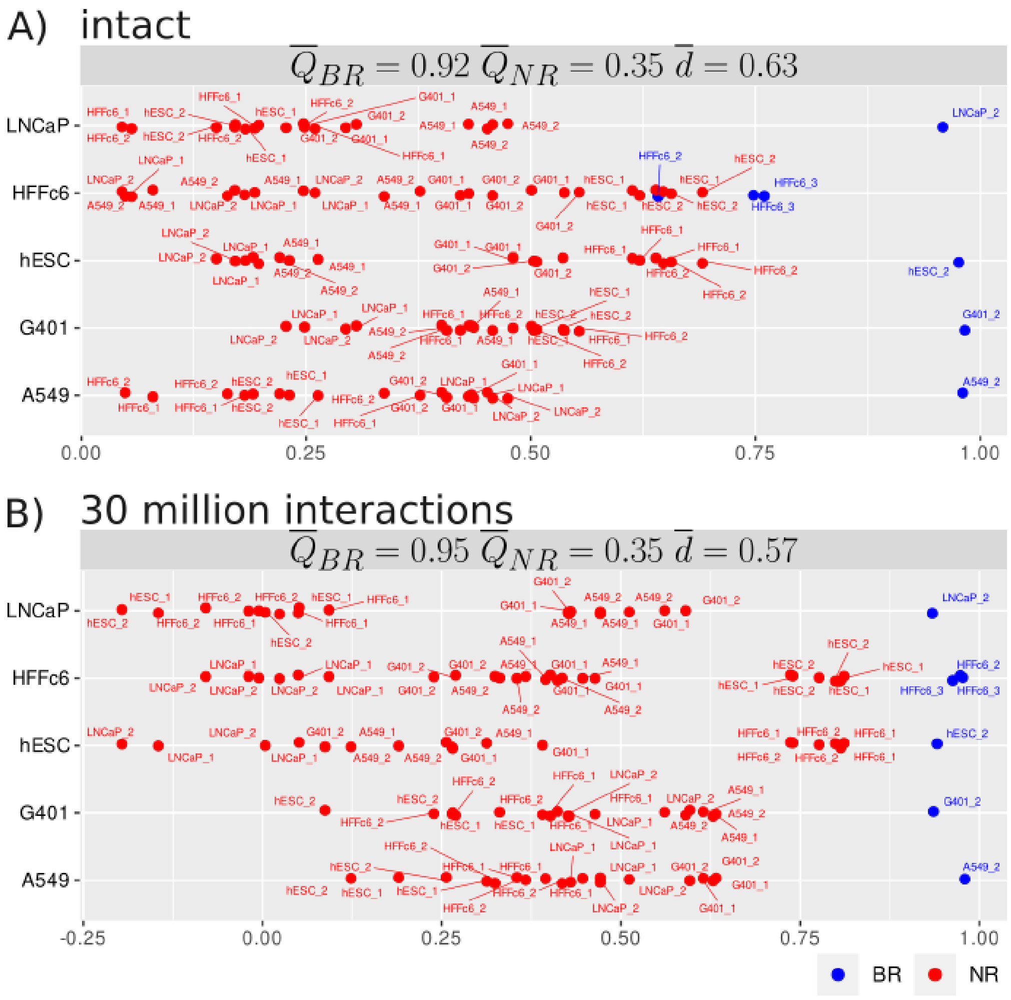


Figure S3: **ENT3C distinguishes biological replicate (BR) contact matrices from non-replicate (NR) contact matrices.** ENT3C similarity scores between BR and NR pairs of 40 kb-binned **(A)** intact contact matrices and **(B)** contact matrices generated from pairs files downsampled to contain 30 million interactions. Each dot represents the similarity score averages across autosomes. ENT3C average similarity scores and separating margins across cell lines ($\overline{Q}_{BR}$, $\overline{Q}_{NR}$ and $\overline{d}$) are indicated in titles (as in Figure 2; Methods). ENT3C parameters: submatrix dimension $n = 300$, window shift $WS = 10$, maximum number of matrices evaluated $WN_{\max} = \infty$.

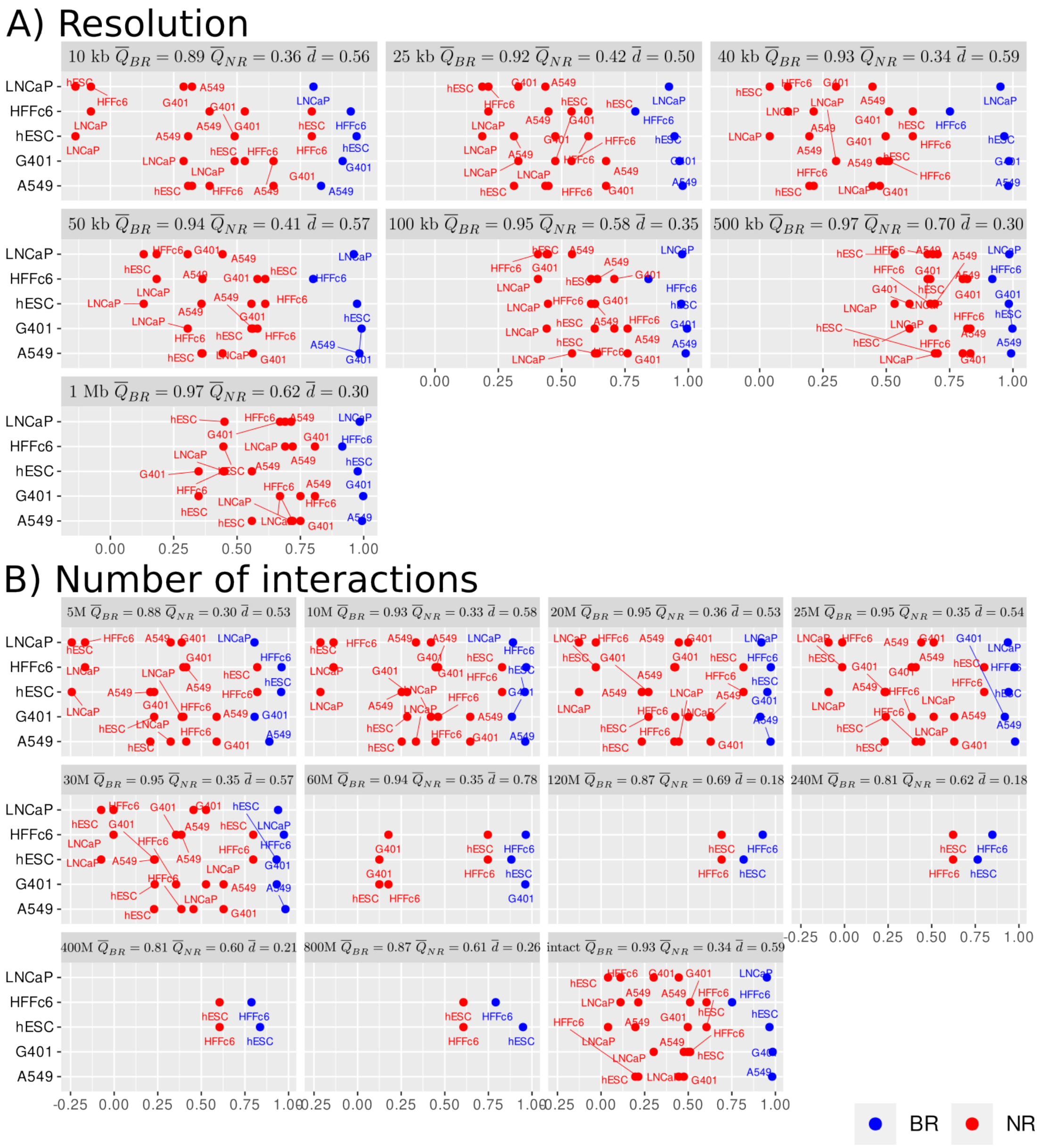


Figure S4: **ENT3C is insensitive to binning resolution and sequencing depth.** Each dot represents ENT3C average similarity scores $Q^{i}_{BR}$ and $Q^{i,j}_{NR}$ (as in Figure 2; Methods) between pairs of **(A)** intact contact matrices binned at 10, 25, 40, 50, 100, 500 and 1000 kb resolutions and **(B)** 40 kb contact matrices generated from pairs files downsampled to 5, 10, 20, 25, 30, 60, 120, 240, 400, 800 million interactions (the last panel indicates intact contact matrices). ENT3C average similarity scores and separating margins across cell lines ($\overline{Q}_{BR}$, $\overline{Q}_{NR}$ and $\overline{d}$) are indicated in titles (Methods). ENT3C parameters: chromosome-split $c = 7$, window shift $WS = 1$, maximum number of matrices evaluated $WN_{\max} = 1000$.

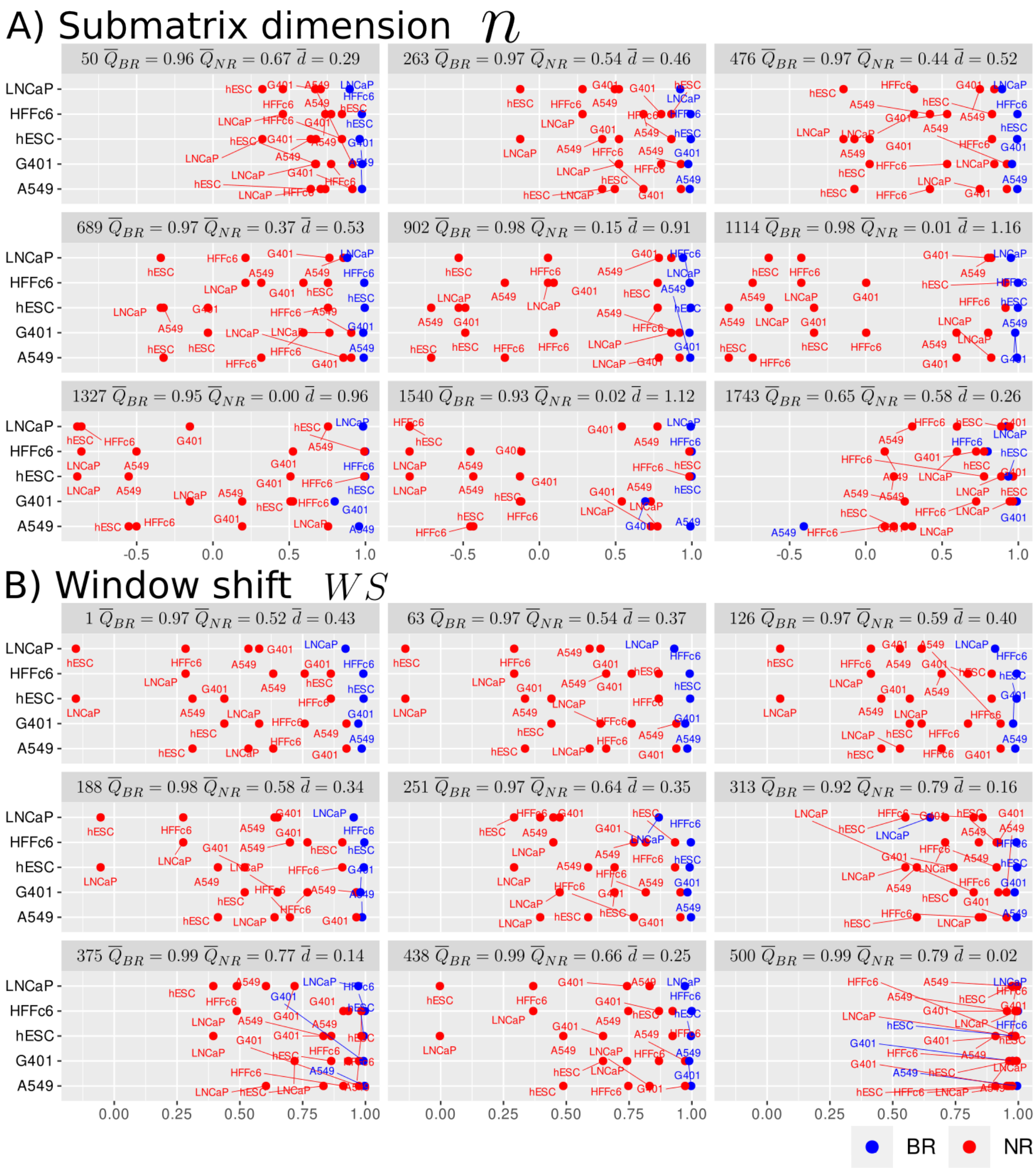


Figure S5: **ENT3C is stable to parameter choice.** Each dot represents ENT3C similarity scores of chromosome 14 $Q^{i}_{BR}$ and $Q^{i,j}_{NR}$ averaged over replicates average similarity scores $Q^{i}_{BR}$ and $Q^{i,j}_{NR}$ (see Figure 2; Methods). Contact matrices for chromosome 14 were generated from pairs files downsampled to 30 million interactions and binned at 40 kb. **(A)** ENT3C's window shift $WS = 1$ and maximum number of matrices evaluated $WN_{\max} = 1000$ were fixed and the submatrix size $n$ was varied between 50 and 1743. **(B)** ENT3C's submatrix dimension $n = 300$ and maximum number of matrices evaluated $WN_{\max} = 1000$ were fixed and the window shift $WS$ was varied between 1 and 500. ENT3C average similarity scores and separating margins across cell lines ($\overline{Q}_{BR}$, $\overline{Q}_{NR}$ and $\overline{d}$) are indicated in titles (Methods).

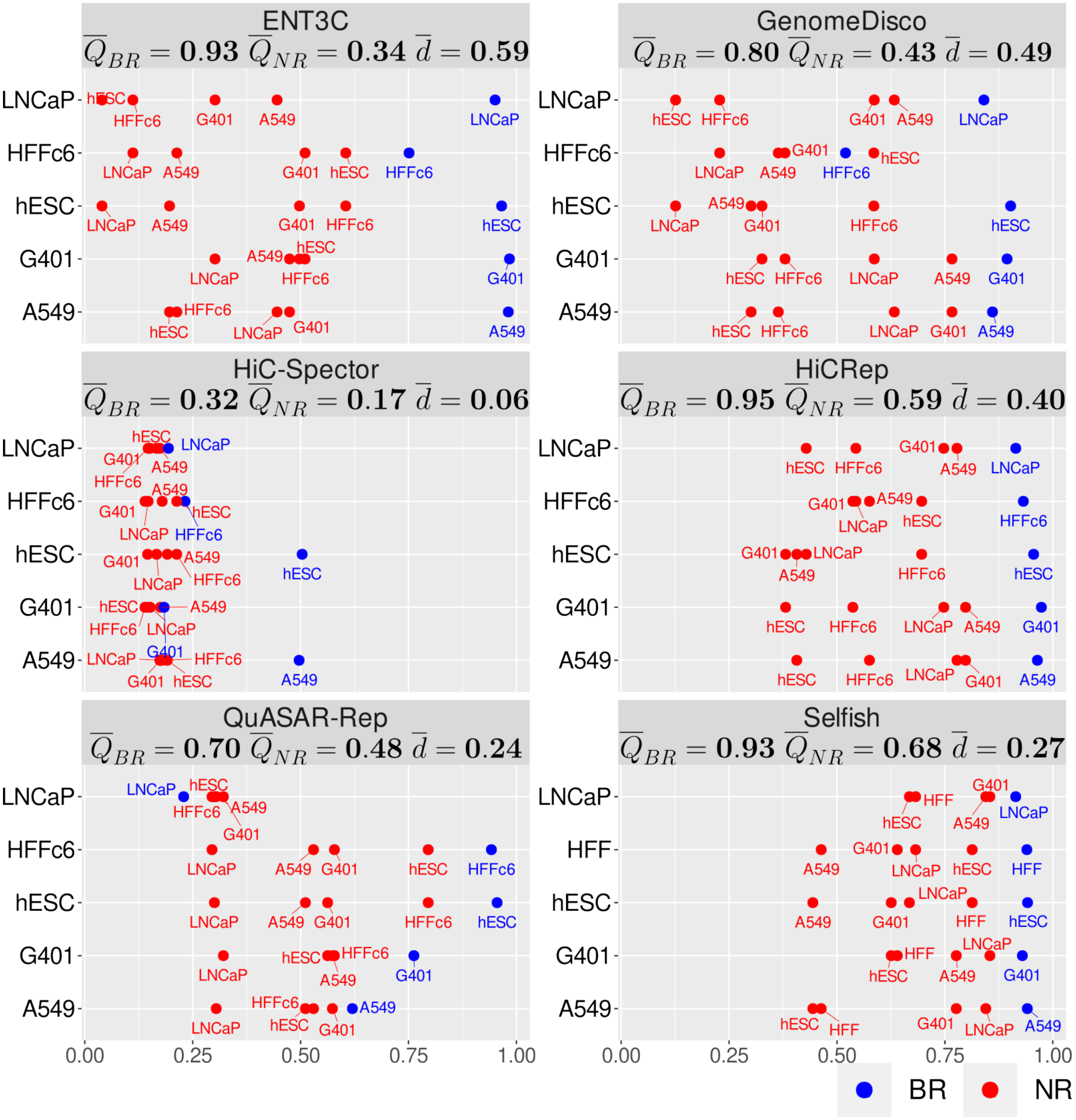


Figure S6: **ENT3C competes well with other methods quantifying Hi-C or micro-C contact matrix similarity.** Each panel represents a Method (ENT3C, GenomeDISCO, HiC-Spector, HiCRep, QuASAR and Selfish) and each dot represents the average similarity scores $Q_{BR}$ and $Q_{NR}$ (as in Figure 2-3; Methods). Intact 40-kb binned contact matrices were used. ENT3C average similarity scores and separating margins across cell lines ($\overline{Q}_{BR}$, $\overline{Q}_{NR}$ and $\overline{d}$) are indicated in titles (Methods). ENT3C parameters: chromosome split $c = 7$, window shift $WS = 1$ and $WN_{\max} = 1000$.

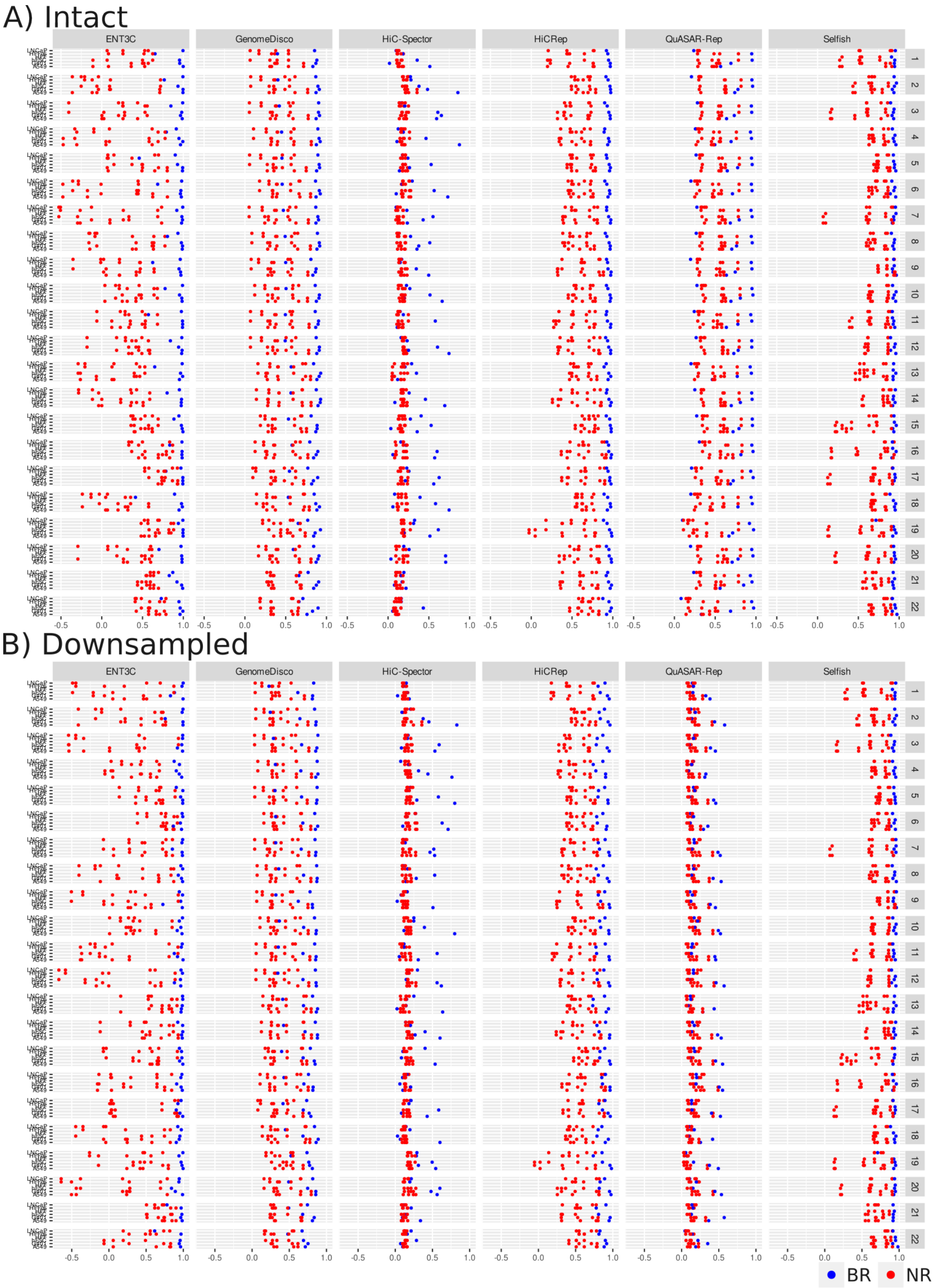


Figure S7: **Contact matrix similarity measures often exhibit chromosomal dependency.** Each dot represents ENT3C similarity scores averaged over replicates between pairs of **(A)** intact contact matrices and **(B)** contact matrices generated from pairs files downsampled to 30 million interactions. ENT3C parameters: chromosome-split $c = 7$, window shift $WS = 1$, maximum number of matrices evaluated $WN_{\max} = 1000$.

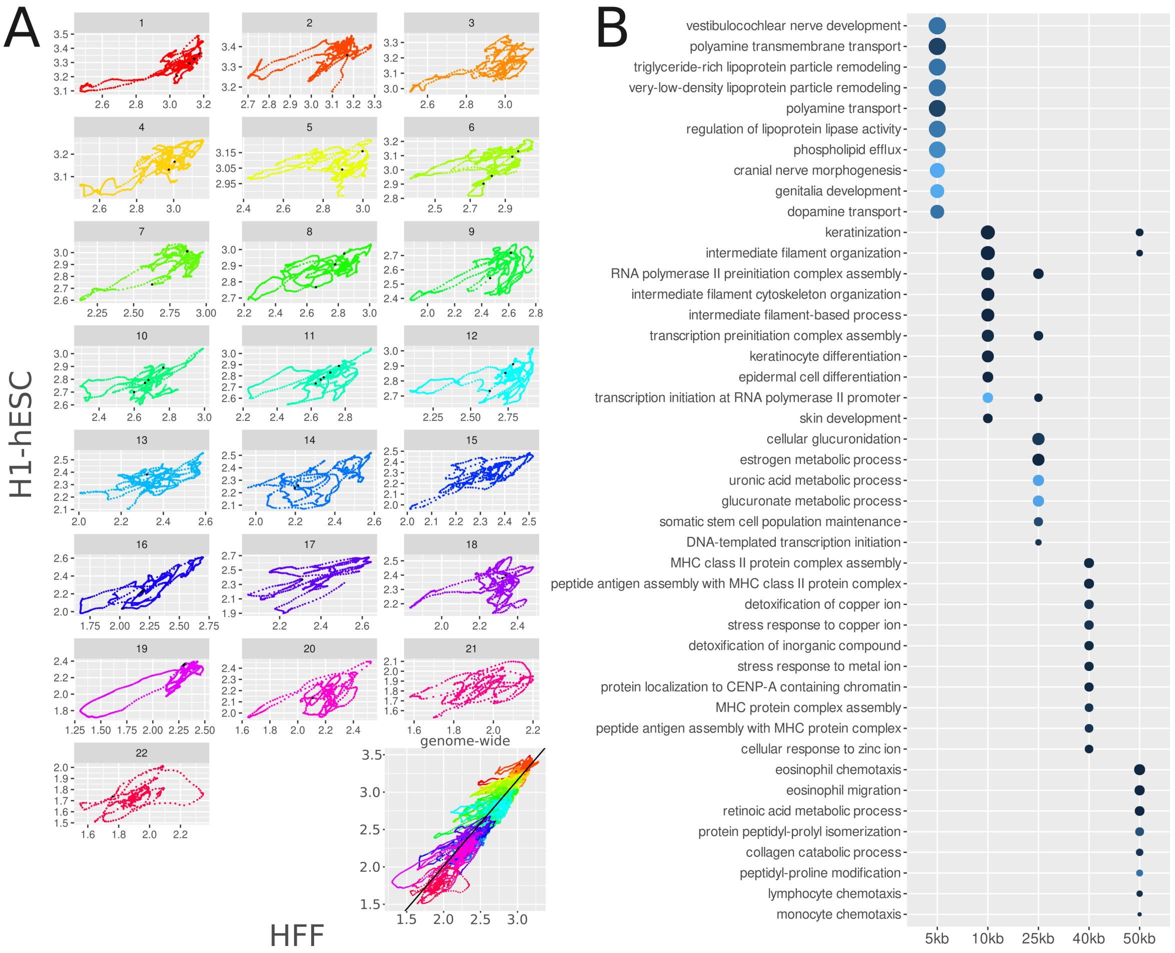


Figure S8: **ENT3C's entropy signals can be used for investigating the biological role of similarly complex regions between two cell lines.** (A) Correlation plots between entropy signals of HFF and H1-hESC 50 kbp contact matrices show interesting patterns. Black points indicate most similar regions. Each chromosome is shown in a different color. In the bottom right genome-wide correlation plot, the genome-wide linear regression line used for determining the most similar regions is shown. (B) Top 10 GO terms arranged by $\log_2$-fold enrichment associated with the genes in the most similar regions at each resolution.

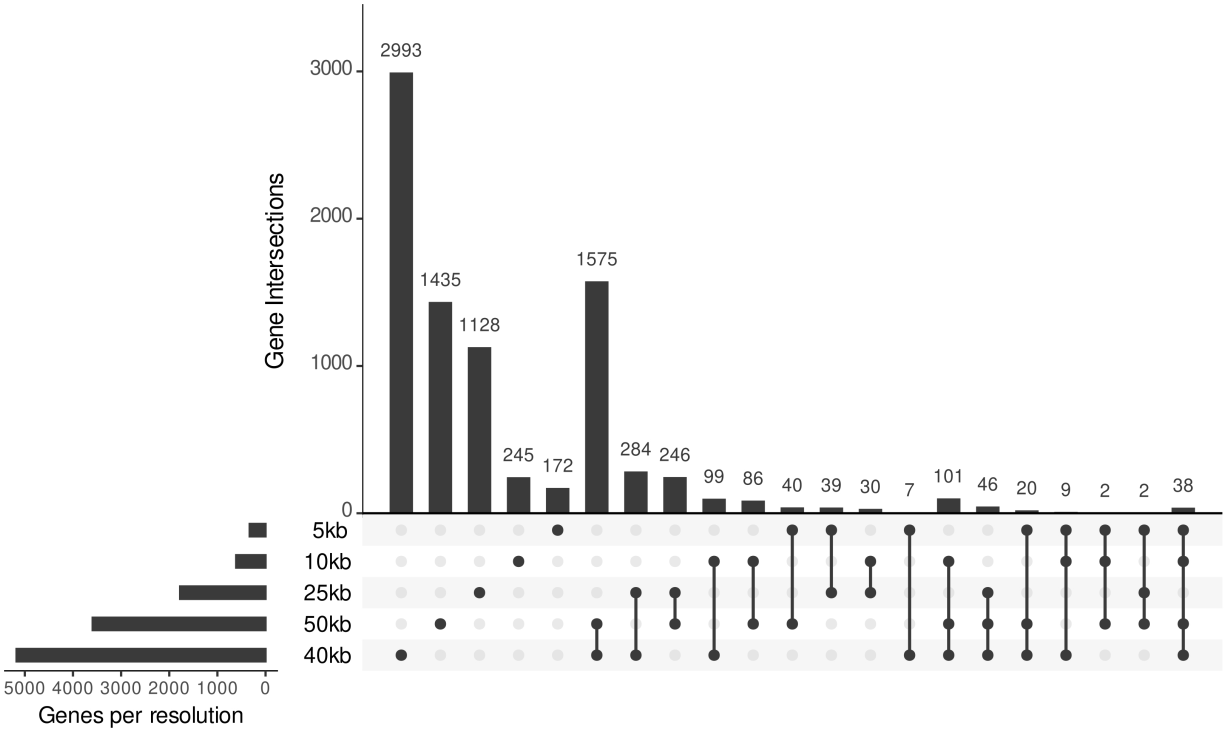


Figure S9: Upset plots of overlapping genes used in GO analysis (Supplementary Fig. S8).

### S1 Supplementary Tables

Table S1: Data sets Hi-C

| Cell line | BR | `BAM` Accession |
|---|---|---|
| G401 | 1 | ENCFF649MAY |
| G401 | 2 | ENCFF758WUD |
| LNCaP | 1 | ENCFF977XHB |
| LNCaP | 2 | ENCFF204XII |
| A549 | 1 | ENCFF867DCM |
| A549 | 2 | ENCFF532XBC |

Table S2: Data sets micro-C

| Cell line | BR | `pairs` Accession |
|---|---|---|
| H1-hESC | 1 | 4DNFING6ZFD, 4DNFIBMG8YA3, 4DNFIMT4PHZ1, 4DNFI8GM4EL9 |
| H1-hESC | 2 | 4DNFIIYUGYBU, 4DNFI89L17XY, 4DNFIXP9MVBU, 4DNFI2YHYAJO, 4DNFIULY29IQ |
| HFFc6 | 1 | 4DNFIN7IIIY6, 4DNFIJZDEIZ3, 4DNFIYBTHGNA, 4DNFIK8UIB5B |
| HFFc6 | 2 | 4DNFIF5F4HRG, 4DNFIK82YRNM, 4DNFIATCW955, 4DNFIZU6ADT1, 4DNFIKWV6BY2 |
| HFFc6 | 3 | 4DNFIFJL4JIH, 4DNFIONHB78N, 4DNFIG1ZOVIM, 4DNFIPKVL9YI, 4DNFIJM966UR, 4DNFIV8JNJB8 |

Table S3: Number of interactions in each sample

| Cell line | BR1 | BR2 | BR3 |
|---|---|---|---|
| H1-hESC | 1255082197 | 153084436 | |
| HFFc6 | 2857647629 | 830616793 | 1423953776 |
| G401 | 100949135 | 101518640 | |
| LNCaP | 53446400 | 35777493 | |
| A549 | 49612977 | 47475253 | |

Table S4: Downsampling of the analyzed datasets

| Nr. Interactions | H1-hESC | HFFc6 | G401 | LNCaP | A549 |
|---|---|---|---|---|---|
| 1 million | 1 | 1 | 1 | 1 | 1 |
| 5 million | 1 | 1 | 1 | 1 | 1 |
| 10 million | 1 | 1 | 1 | 1 | 1 |
| 20 million | 1 | 1 | 1 | 1 | 1 |
| 25 million | 1 | 1 | 1 | 1 | 1 |
| 30 million | 1 | 1 | 1 | 1 | 1 |
| 60 million | 1 | 1 | 1 | 0 | 0 |
| 120 million | 1 | 1 | 0 | 0 | 0 |
| 240 million | 1 | 1 | 0 | 0 | 0 |
| 400 million | 1 | 1 | 0 | 0 | 0 |
| 800 million | 1 | 1 | 0 | 0 | 0 |
